# Inborn errors E778K and G908R in NOD2 gene increase risk of nontuberculous mycobacterial infection: a computational study

**DOI:** 10.1101/2020.12.25.424387

**Authors:** Shamila D. Alipoor, Mehdi Mirsaeidi

## Abstract

**Background:** The innate immune system has a critical role in the early detection of pathogens, mainly by relying on pattern-recognition receptor (PRR) signaling molecules. Nucleotide-binding oligomerization domain 2 (NOD2) is a cytoplasmic sensor for recognition of invading molecules and danger signals inside the cells. NOD2’s functions are critical; polymorphisms of its encoding gene are associated with several immune pathological conditions. We recently reported that missense E778K and G908R variants of NOD2 gene are associated with recurrent pulmonary nontuberculous mycobacterial infections

**Methods:** This is an *in-silico* analysis of NOD2 gene using SNPs functionality analyses, post-translational modification site prediction and network analysis.

**Results:** Our analysis revealed that these damaging mutations affect the structural properties and function and ligand binding in the mutant receptor.

**Conclusion:** The consequence of these mutations may also impress downstream processing and receptor crosstalk with other immune molecules and therefore increase susceptibility to infectious disease.

## 1. Introduction

It has been suggested that individual genetic composition plays an important role in susceptibility to pulmonary nontuberculous mycobacterial (PNTM) disease^1–3^. The innate immune system provides the first line of defense against the danger signals and relies mainly on the pathogen recognition receptors (PRRs). PRRs are the main immune system players that can detect the molecules that frequently are found in pathogens or released by damaged cells (respectively known as Pathogen/ damage associated molecular patterns -PAMPs or DAMPs)^4,5^.

The PRRs can be categorized into four distinct functional groups: (1) Toll-like receptors (TLRs), (2) retinoic acid–inducible gene (RIG)-I-like receptors (RLRs), (3) C-type lectin receptors (CLRs) and (4) Nucleotide-binding oligomerization domain-like receptors (NLR)^6^. Nucleotide-binding oligomerization domain-containing protein 2 (NOD2) was NLR discovered in 2000, after the discovery of NOD1 in 1999 ^6,7^. The NOD2 gene is situated on chromosome 16 and encodes the NOD2 protein^8^. The role of NOD2 in macrophages and bronchial epithelial cells exposed to mycobacteria has been summarized in Figure 1 and 2. It detects a fragment of bacterial cell wall named peptidoglycan muramyl dipeptide (MDP) and subsequently activates the signaling pathways leading to proinflammatory cytokine production^9,10^.

**Figure 1.**
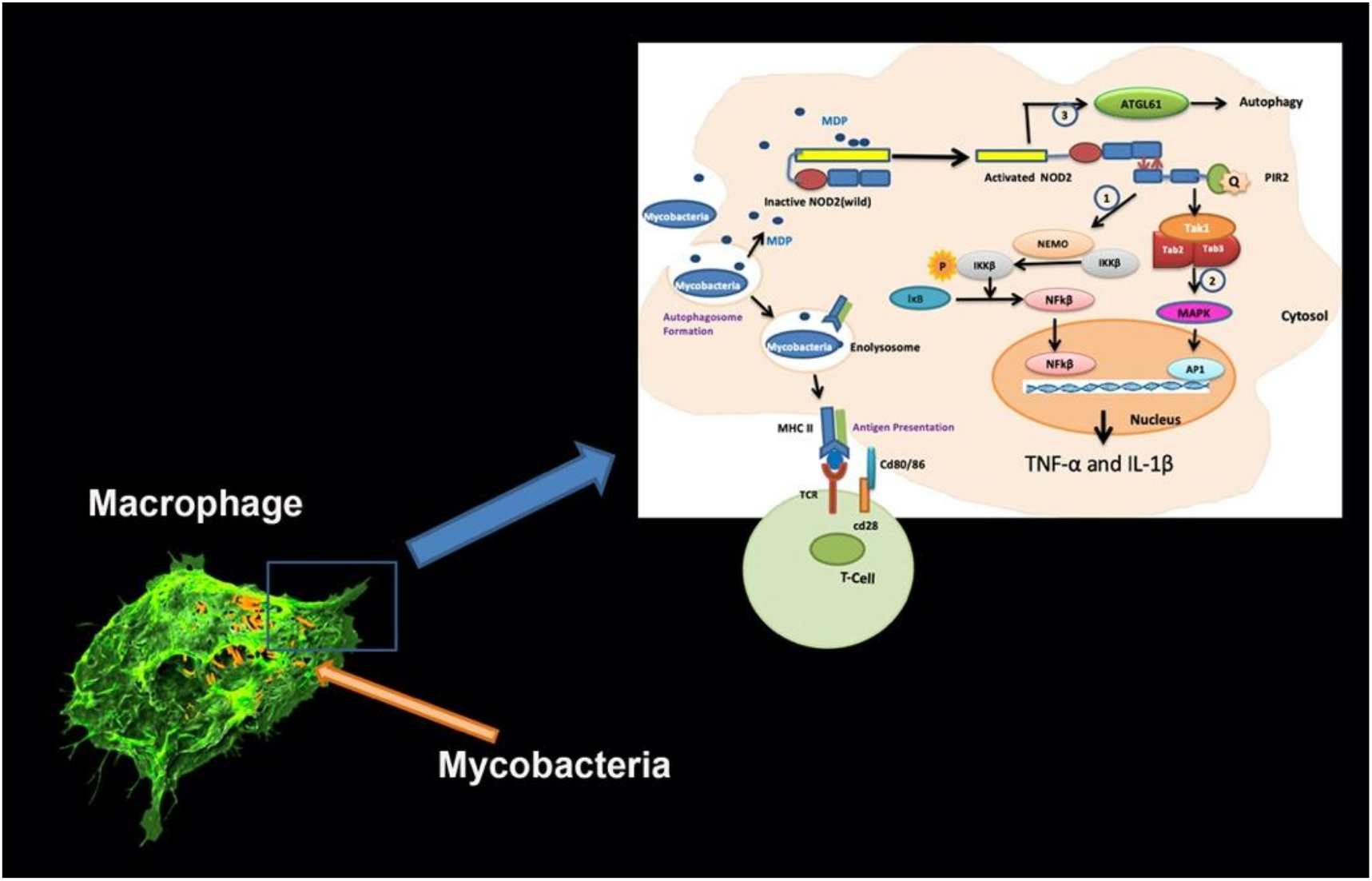
Summary of the role of NOD2 in response to mycobacterial infection in macrophages^19–21^. Upon ligand binding and receptor activation, the C-terminal LRR domain of NOD2 undergoes a conformational change and exposes the CARD domain which allows it to interact and oligomerize with the CARD domain in the adaptor molecule RIP2 (receptor-interacting protein 2) through a homophilic CARD-CARD interaction. Upon oligomerization, activated PIR2 applies “lysine 63 (K63)-linked polyubiquitination” at lysine 209, located at the kinase domain of the molecule. This ubiquitination promotes recruitment of TAK1 and NEMO (the NF-kB essential modulators). The activation of TAK1 and NEMO promotes phosphorylation of the IKKβ, which is a key kinase in the NF-κB signaling pathway. Phosphorylated IKKβ degrades IκB, and subsequently activates NF-κB family transcription factors. In addition, this path also activates the mitogen-activated protein kinases (MAPKs), ultimately triggering transcription factor activator protein 1 (AP1). The NF-κB and MAPK pathways together stimulate the expression of inflammatory cytokines. Activated NOD2 also may trigger an autophagic pathway by recruiting ATG16L1.

**Figure 2.**
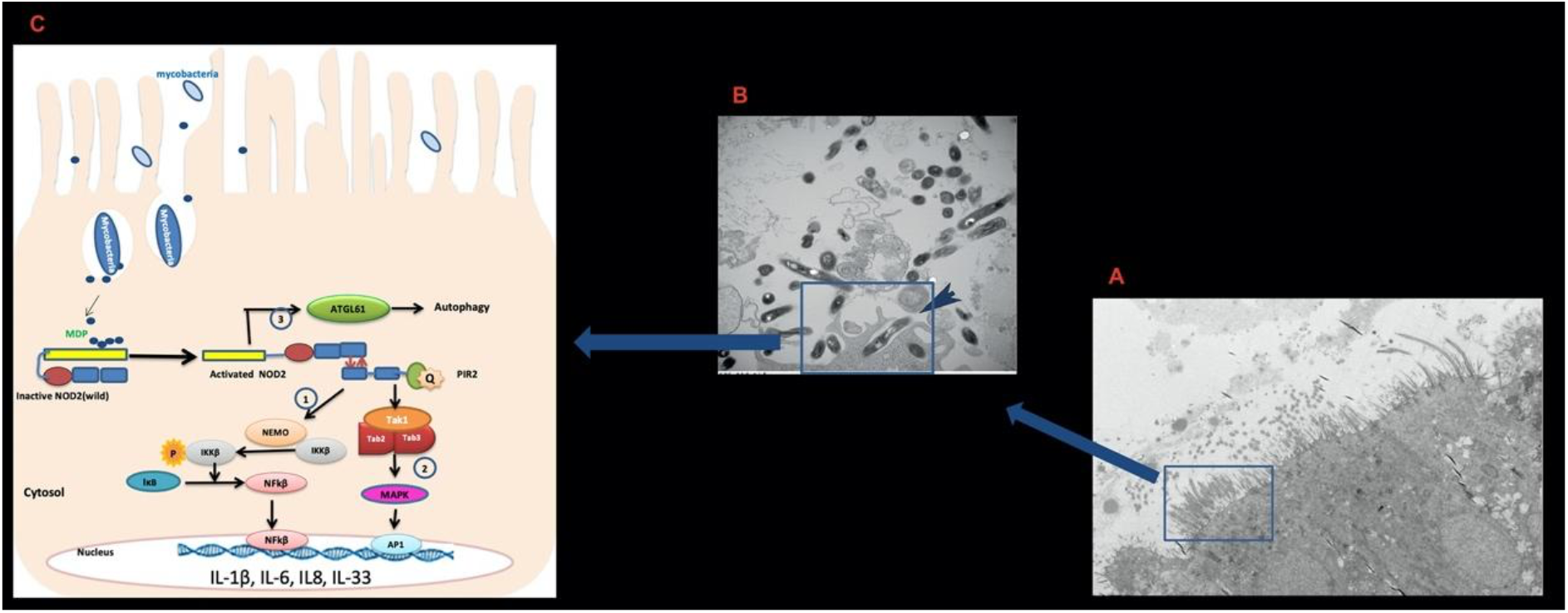
Summary of the role of NOD2 in response to mycobacterial infection in pulmonary epithelial cells^22,23^. (A) Bronchial epithelial cells with cilia, (B) Engulfment of mycobacteria in epithelial cells, C. Subcellular NOD2 model of action.

The leucine-rich repeats (LRR) domain of NOD2 senses molecular determinants from a structurally diverse set of bacterial, fungal, parasitic and viral-derived components. The LRR domain of NOD2 covers amino acids 744 to 1020 and consists of repeated LRR units. This domain contains 9 helixes and 11 sheets, which forms a typical horseshoe-like structure in a single curvature^11^.

The C-Terminal LRR domain is responsible for the detection of PAMPs and DAMPs and also negatively regulates protein activity. Upon ligand binding, the LRR domain undergoes a conformational change and exposes the CARD domain to interaction with downstream molecules^12^. Generally, upon ligand binding, the closed form (inactive) of the protein changes to the open (active) form with WHD–HD2–LRR rotation relative to the NBD–HD1 modules^11^.

It has been shown that the polymorphisms in the NOD2 gene are associated with the occurrence of some infectious diseases and granulomatous inflammation, including mycobacterium tuberculosis^13^, Crohn’s disease ^14^, and Blau’s syndrome ^15–17^.

We recently reported 3 patients with inborne errors in the LRR region of the NOD2 gene (E778K and G908R) with recurrent PNTM ^18^; however, the underlying mechanism of mutated NOD2 for increasing susceptibility to infections, including NTM, is not yet understood.

We aimed to assess the effect of missense E778K and G908R mutations on the stability, structural properties, and interdomain and interatomic interactions of NOD2. Given that E778K and G908R are located in the LRR domain, we hypothesized that these inborne errors alter host sub-cell signaling by stopping the antigen presenting process.

## 2. Material and Methods

### 2.1. Data mining

The Fasta format of the NOD2 sequence was extracted from Uniprot^24^. The secondary structures of the wild and mutant proteins were extracted from the pdb files by DSSP^25^. Solvent accessibility of the residues in the structures of the proteins was extracted from the pdb file using Getarea^26,27^.

### 2.2. Characterization of functionality of SNPs

Evaluation of the Impact of E778K and G908R was performed using Sequence Homology Tools. In order to identify the effects of amino acid substitution on the phenotypic and functional properties of the receptor molecule, we used the Provean analysis tool (http://provean.jcvi.org/index.php). Provean predicts tolerated and deleterious SNPs based on the multiple alignment information for every position of the sequence^28^.

In this work, the fasta SEQ obtained from NCBI and the mutated positions were submitted as a query to Provean for homology searching. The substitutions at each position equal to or less than a threshold of −2.5 indicates an intolerant or deleterious variant.

### 2.3. Prediction of the functional consequence of SNPs by PolyPhen

To determine the possible consequence of amino acid substitutions on the function of the NOD2 receptor, we used PolyPhen-2 (http://coot.embl.de/PolyPhen)^4^. For this analysis, the amino acid sequence of NOD2 wild type in fasta format was used as input query and the position and substitution of each of the two SNPs were submitted to PolyPhen.

PolyPhen works by a set of supervised learning algorithms that act based on the straightforward empirical rules applied to the input sequence in order to characterize the substitutions [5].

PolyPhen first calculates a score for the position-specific independent counts (PSIC) based on the multiple alignment for each of the two amino acid residues (original and mutant), and then computes the PSIC scores as the difference of the two residues. The mutations are characterized based on the cutoff PSIC index as probably damaging (PSIC score > 0.85), possibly damaging (PSIC score > 0.15), or benign. The functional impact of a particular amino acid substitution with a higher PSIC score is more likely to be harmful.

### 2.4. Characterization of Functional nsSNPs by PANTHER

PANTHER (http://www.pantherdb.org) was used to characterize the two SNPs based on an HMM statistical model and evolutionary relationship ^29^.

Functional Analysis through Hidden Markov Model (fathmm)^30^ using PANTHER-cSNP scoring tool was performed to predict the functional and phenotypic consequences of missense variants by combining sequence conservation within hidden Markov models (HMMs)^29^.

The amino acid substitution and protein fasta sequence was used as inputs of cSNP scoring tool for PANTHER, and a score was computed by the tool, based on the position-specific evolutionary preservation (PSEP) score. The chosen thresholds are “probably damaging” (time > 450my), “possibly damaging” (450my > time > 200my), and “probably benign” (time < 200my).

### 2.5. Identification of functional SNPs in conserved regions by ConSurf

To identify the evolutionary conservation of the amino acids in the protein sequence, we employed ConSurf, which analyzes the phylogenetic relationships between homologous sequences^31^.

### 2.6. Determination of the SNPs’ structural effects on protein stability and flexibility

To analyze how the mutation affects the structure and stability of the protein, we used I-Mutant^32^. I-Mutant is a web server that acts based on a support vector machine that predicts the stability of a mutant protein using a ProTerm-derived dataset. We submitted the SNPs of the NOD2 sequence in fasta format as input. We also performed further analysis using DynaMut ^33^ to assess the flexibility and interatomic interactions in wild and mutant proteins.

### 2.7. Prediction of ligand binding sites

The prediction of ligand binding sites on the wild and mutant models was performed by FTSite ^15,34^.

### 2.8. Analysis of structural specificity of functional SNPs (HOPE)

To understand the effect of the SNPs on the structure of the protein we used HOPE ^35^. HOPE is a web-server tool that identifies the structural effects of point mutations in a protein sequence. We used the NOD2 wild type sequence and the 2 SNPs individually as the input.

### 2.9. Prediction of post-translational modification site

Search for the putative PTM sites in the human NOD2 protein was performed using NetPhos ^36^. The NetPhos 3.1 server predicts the Serine, Threonine, and Tyrosine phosphorylation sites in the protein. For searching ubiquitination sites and peptide cleavage sites we used BDM-PUB^37^ and ProP 1.0 ^38^ respectively.

### 2.10. Network analysis

For further analysis of the effects of these SNPs, we analyzed the interaction of the mutant NOD2 with other downstream proteins.

Protein–protein interaction (PPI) networks were constructed and analyzed by STRING and Cytoscape v. 3.4.0 [30]. Briefly, NOD2 was introduced to the software as the seed protein, and PPI networks were constructed using STRING ^39^ and visualized and analyzed in Cytoscape. STRING provides information on both the experimental and the predicted interactions based on gene neighborhood, gene co-occurrence, gene fusions, gene co-expression, and literature mining. Highly interconnected regions (clusters) and their related pathways were extracted.

### 2.11. Differential contact map analysis

For finding the differences between the contact maps of wild and mutant proteins, we used a method described by Lyer et al ^40^. Briefly, we first calculated a differential contact map (DCM) by subtracting one contact map from the other. Then, we filtered this map and removed any residue pairs that showed less than (or equal to) 5Å difference in inter-residue distance in the two conformations. This was done in order to consider only the residues that could be modulating large scale conformation changes and to exclude contact changes seen as a result of small local fluctuations in the structure^40^.

## 3. Results

### 3.1. Characterization and functionality of missense E778K and G908R variants

Functionality of missense E778K and G908R variants was evaluated by predicting which substitution of the amino acids are critical for NOD2 function by using in silico sequence homology Tools, including Protein Variation Effect Analyzer (Provean), Polymorphism Phenotyping-2 (PolyPhen-2), (Protein Analysis Through Evolutionary Relationships) (PANTHER), and conSurf computational tools (Table 1).

**Table 1:** Characterization of functionality of SNPs using Sequence Homology Tools

|  | E778K |  | G908R |  |
| --- | --- | --- | --- | --- |
|  | Score |  | Score |  |
| Provean (Tolerated index= -2.5) | -2.57 | Damaging | -5.822 | Damaging |
| Polyphen-2 | 0.98 | Probably Damaging | 1.0 | Probably Damaging |
| PANTHER | 455my | Damaging | 457my | Damaging |
| HOPE |  | Damaging |  | Damaging |

Provean demonstrated that both E778K and G908R were deleterious (score of −2.57 and −5.822 respectively with cutoff=−2.5) (Table 1).

To further understand the effects of amino acid substitution on the structure and function of NOD2, the PolyPhen-2 score was calculated for E778K and G908R. PolyPhen-2 was scored as 0.98–1.00. Variants with these ranges of scores are more confidently predicted to be damaging to NOD2 structure and functionality.

PANTHER-PSEP analysis showed a high score (PSEP score> 450 million years) for E778K and G908R, suggesting highly specific evolutionary preservations for both positions causing damage to the protein’s functionality. Evolutionary conservation analysis was also performed by the ConSurf web tool. It characterized structural and functional residues, using evolutionary conservation and solvent accessibility. ConSurf showed that E778K is an exposed and functional position whereas G908R is a buried and structural position. Both positions are highly conserved (Figure 3) so substitutions would be high risk and would damage receptor function.

**Figure 3.**
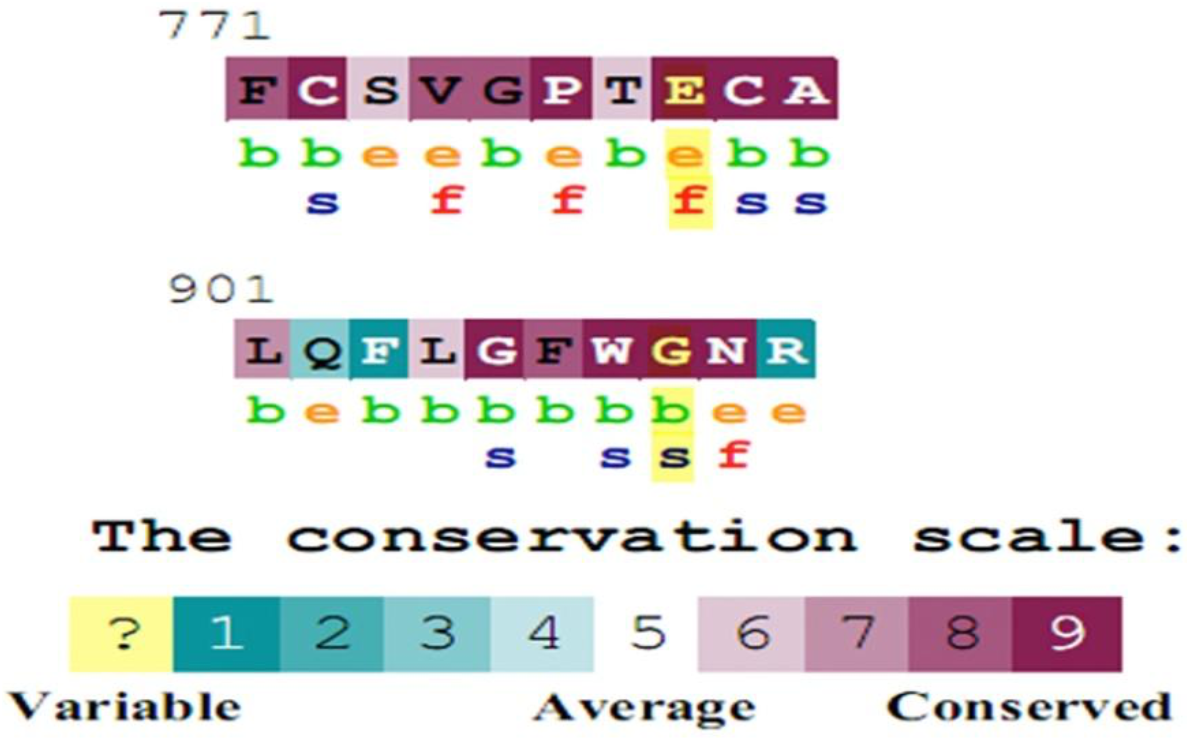
Evolutionary conservancy of NOD2; mutated positions produced by Consurf (e: An exposed residue according to the neural-network algorithm. b: A buried residue according to the neural-network algorithm. f: A predicted functional residue (highly conserved and exposed). s: A predicted structural residue (highly conserved and buried)). Results are provided by ConSurf.

### 3.2. Structure analysis of wild type and mutant models

Homology models of the human NOD2 in wild and mutant forms (chain A) were built by Swiss models using the 5irm (Crystal structure of rabbit NOD2) as a template.

Structural similarities between native and mutant models were investigated based on TM-score and RMSD scores using Bio3d and TM-align tool. The total TM-score and the RMSD between wild and mutant structures were 0.99 and 0.15 in the case of E778K and 0.99 and 0.38 in the case of G908R respectively; that indicated that the two structures are in almost the same fold.

### 3.3. The effect of the mutations on the NOD2 structure

The Project HOPE analysis revealed that the mutant residues in both SNPs are different in size and hydrophobicity properties compared to the wild type residues, and these variations can disrupt the interatomic interactions, including the H-bond and ionic interactions; salt bridge and H-bond with the adjacent molecules (Table 1; Figure 4 and 5).

**Figure 4.**
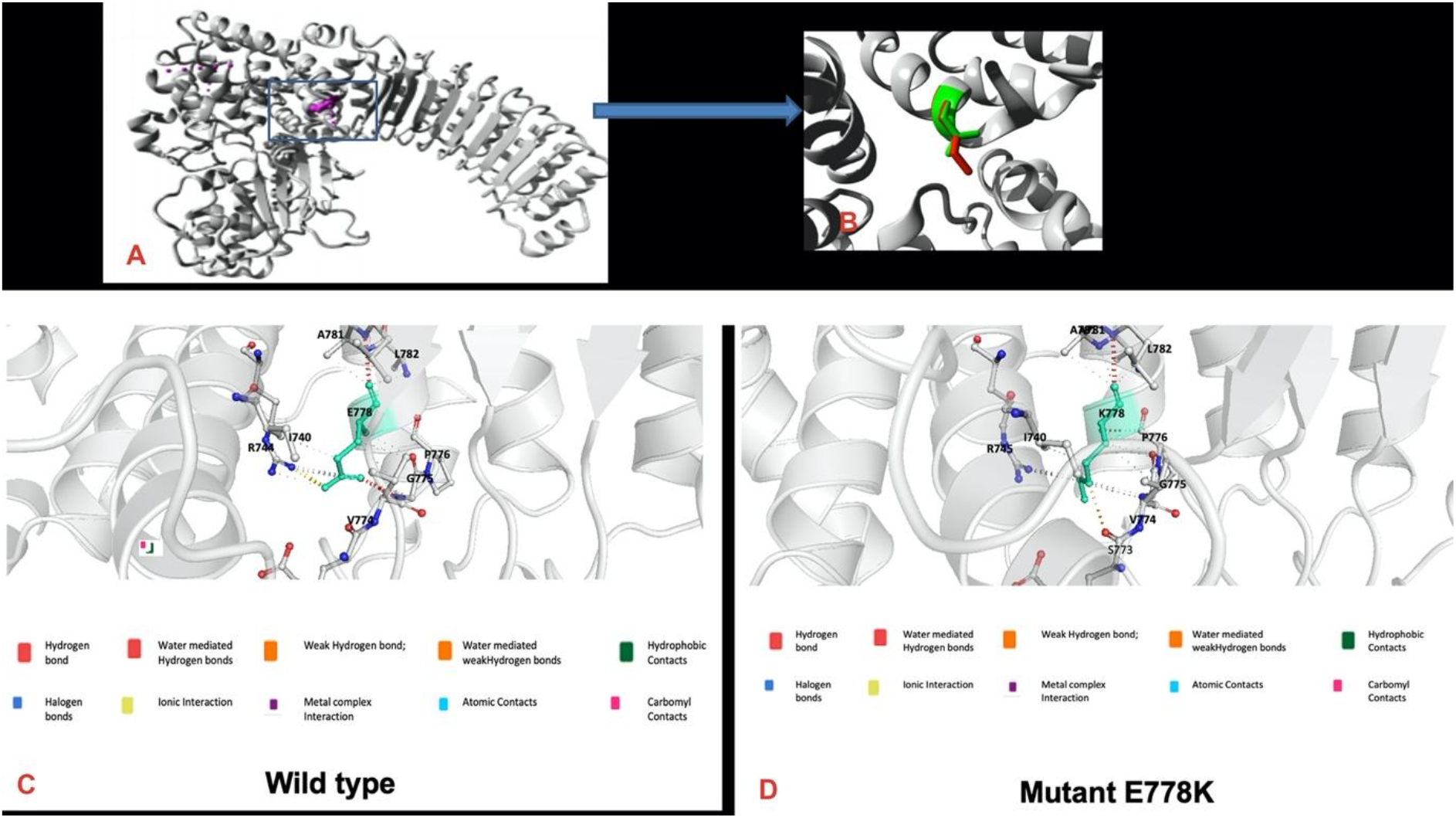
Structural change mutation E>K illustrated by HOPE project: (A) Overview of the protein in ribbon-presentation. The mutated residue E778K is colored and shown as small balls. (B) The protein is colored grey; the side chains of wild-type and the mutant residue are shown green and red respectively. (C, D) Prediction of interatomic interaction by DynaMut in E778K. (C) Wild-type and (D) mutant residues are colored in light-green and are also represented as sticks alongside the surrounding residues which are involved in any type of interactions.

**Figure 5.**
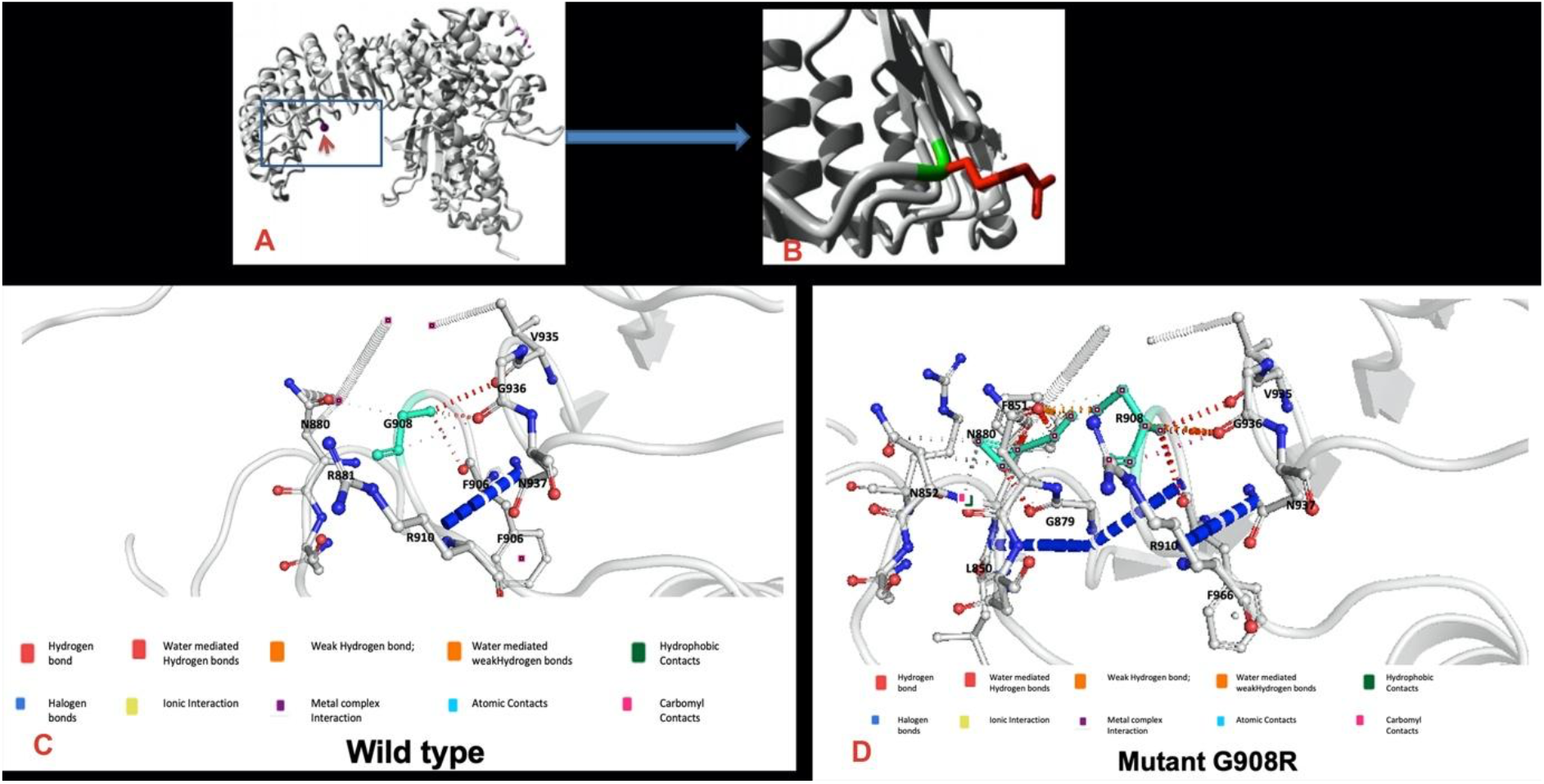
(A, B) Structural change mutation G908R illustrated by HOPE project: (A) Overview of the protein in ribbon-presentation. The mutated residue is colored and shown as small balls. (B)The protein is colored grey; the side chains of wild-type and the mutant residue are shown green and red respectively. (C, D) Prediction of interatomic interaction by DynaMut in G908R. (C) Wild-type and (D) mutant residues are colored in light-green and are also represented as sticks alongside the surrounding residues which are involved in any type of interactions.

E778K, is located on the α2 helix in LRR7. Glu (E) is an acidic while Lys (K) is a basic amino acid. K is bigger than Glutamic acid. The wild-type residue charge is negative while the mutant residue has a positive charge. This can cause repulsion with other residues in the protein or ligands^35^ (Figure 4).

The prediction of interatomic interactions with DynaMut revealed that the mutation could significantly change interaction between residues in the mutant proteins (Figure 4).

Additionally, missense 3D tools predicted that E778K substitution leads to the expansion of a cavity (a pocket on the protein surface) volume by 9.504 Å^3.

G908R is located on a turn between the α6 and α7 helixes in the LRR domain and is not solvent accessible (Figure 5). Glycine (G) is a small and nonpolar amino acid, while Arginine (R) is considered to be a basic amino acid with a larger side chain. The G residue charge is neutral while the mutant residue has positive charge. Furthermore, G is more hydrophobic than R. G is considered to be the most flexible of all the residues necessary for the protein’s function. Mutation of this G can abolish this function and also might disturb the LRR repeat and consequently any function this repeat might have. The mutation causes repulsion of ligands or other residues with the same charge.

DynaMut prediction of interatomic interactions revealed that changing G to R significantly changes the interaction of new residue with the adjacent residues in the mutant proteins. Additionally, missense 3D tools predicted that G908R leads to the expansion of a volume cavity on the protein surface by 27.432 Å.

### 3.4. Determination of the SNPs’ effects on protein stability and flexibility

To predict the effects of SNPs on the stability of the NOD2 receptor based on the reliability index (RI) and the free energy change values derived by measuring Delta Delta G (DDG), we used the I-Mutant tool (https://folding.biofold.org/i-mutant/i-mutant2.0.html). The results indicated that both SNPs decreased stability of the protein, as shown in Table 2.

**Table 2:** Effect of SNPs on protein stability predicted by I-MUTANT 2.0

|  | Stability | RI | DDG-Free energy change value (kcal/mol) |
| --- | --- | --- | --- |
| E778K | Decreased | 7 | -0.96 |
| G908R | Decreased | 6 | -1.23 |

DG(NewProtein)-DG(WildType) in Kcal/mol; DDG<0: Decrease Stability; DDG>0: Increase
Stability

We also compared the flexibility of wild and mutated proteins based on vibrational entropy changes using Dynamut. In the case of E778K, the result indicated that mutation E to K increases total molecular flexibility (∆∆SVib ENCoM: 0.133 kcal.mol-1.K-1) and increases protein flexibility in the LRR domain. However, this mutation increased rigidity in the HD1, HD2, and WHD domains of the NOD2 structure (Figure 6).

**Figure 6.**
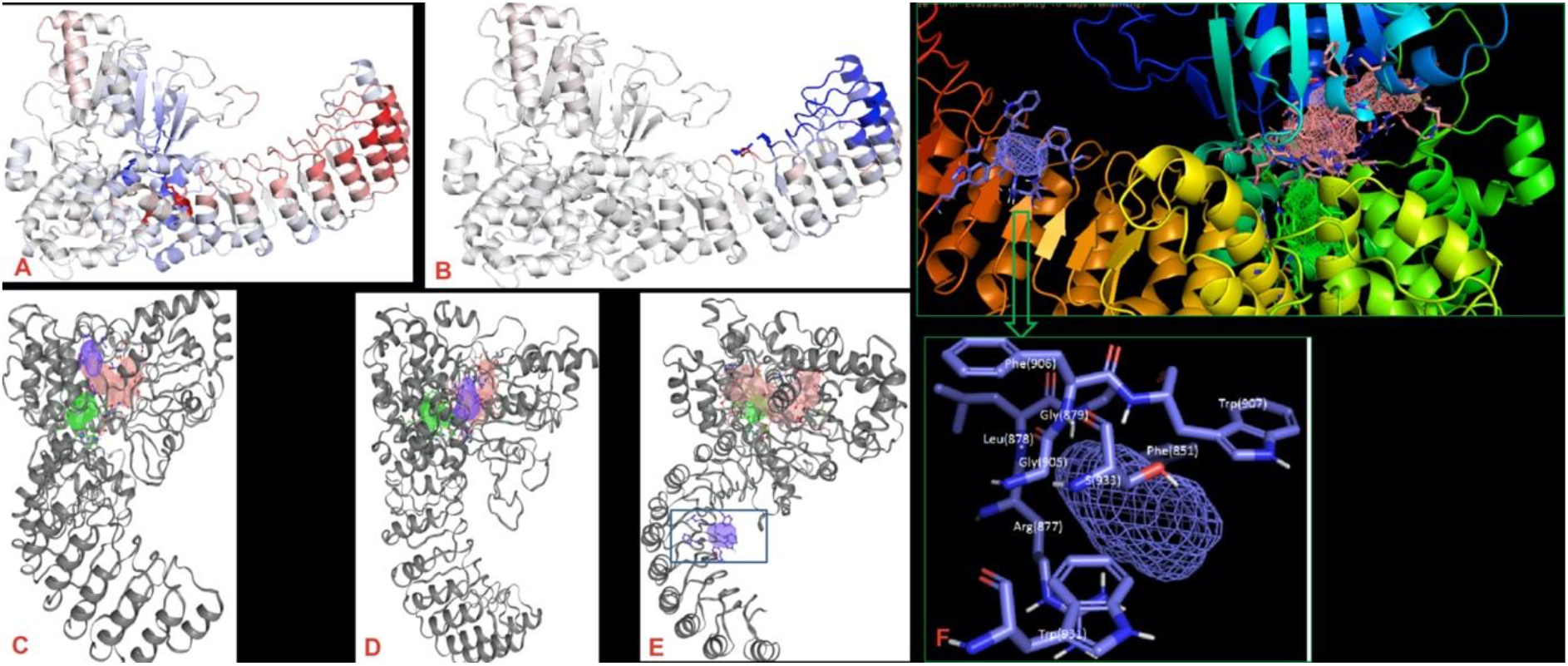
(A) E778K and (B) G908R Amino acids colored according to the vibrational entropy change upon mutation. Blue represents a rigidification of the structure and Red a gain in flexibility. Image is illustrated by Dynamut. (C, D, E) Prediction of interatomic interaction in wild (C), E778K (D), and G908R (E) by DynaMut. Wild-type and mutant residues are colored in light-green and are also represented as sticks alongside the surrounding residues which are involved in any type of interactions. Structures are illustrated by FTSite. Binding site positions are colored in the following order: Site 1 (pink), site 2 (green), and site 3 (blue). (F) Predicted binding sites in the mutant G908R receptor; Site 3 ligand binding site in G908R mutant protein. The region in blue contains phe851; gly 905; phe906, trp907, arg910, arg877; trp931 that form the site 3 binding site.

For G908R, mutation reduces protein stability (ΔΔG: −0.061 kcal/mol) and also decreases total molecular flexibility (ΔΔSVib ENCoM: −0.471 kcal.mol-1.K-1) as shown in Figure 6.

These results indicate that E778K and G908R interfere with conformational change of receptor in response to Muramyl dipeptide of mycobacteria.

### 3.5. Prediction of NOD2 Molecular Binding Sites

FTSite was used for identifying the ligand binding sites for wild and mutant proteins. The results showed three binding sites in the NOD2 structure (Table 3). Site 1 and 2 were similar in wild and mutant structures, but a significant change was observed in binding site 3 of the G908R mutant. In wild and E778K structures, binding site 3 mainly contains 237(L); 242-3(GA); 251(I); 307(T); 455(Y); 486(P); 490(W), while in the G908R mutant, this site includes 851(F); 877-9(RLG); 905-7(GFW); 931-3(W-S) (Table 3) (Figure 6).

**Table 3:** Ligand binding sites in Wild and mutant receptor NOD2 predicted by FTsite

|  | Wild | Mutant 778 | Mutant 908 |
| --- | --- | --- | --- |
| **Site 1** | EAGSGKST(300-306)<br>SCRQ(322-6)<br>DGFDE(379-383)<br>424;489;490;586;600 | EAGSGKST(300-306)<br>SCRQ(322-6)<br>DGFDE(379-383)<br>424;489;490;586;600 | EAGSGKST(300-306)<br>SCRQ(322-6)<br>DGFDE(379-383)<br>424;489;490;586;600 |
| **Site 2** | 301<br>(L—VF)485<br>(TT--Y)510-514<br>T605<br>(I --AF)673-677<br>(W--R---E)741-748 | 301<br>(L—VF)485<br>(TT--Y)510-514<br>T605<br>(I --AF)673-677<br>(W--R---E)741-748 | 301<br>(L—VF)485<br>(TT--Y)510-514<br>T605<br>(I --AF)673-677<br>(W--R---E)741-748 |
| **Site 3** | 237(L);242-3(GA);251(I);307(T);<br>455(Y); 486(P); 490(W) | 237(L);242-3(GA);251(I);307(T);<br>455(Y); 486(P); 490(W) | 877-9(RLG);905-8(GFW);931-3(W-S)L-T(237) |

### 3.6. Contact map analysis

For further analysis of interdomain interactions, the differential contact map of the residue in the wild and mutated form was accessed. For this analysis, a homology model was built for the NOD2 structures with both mutations and was compared to the wild type protein. The analysis showed that in the mutant model, the residues’ contacts in the CARD domain are changed upon introduction of the mutations in the LRR domain (Figure 7).

**Figure 7.**
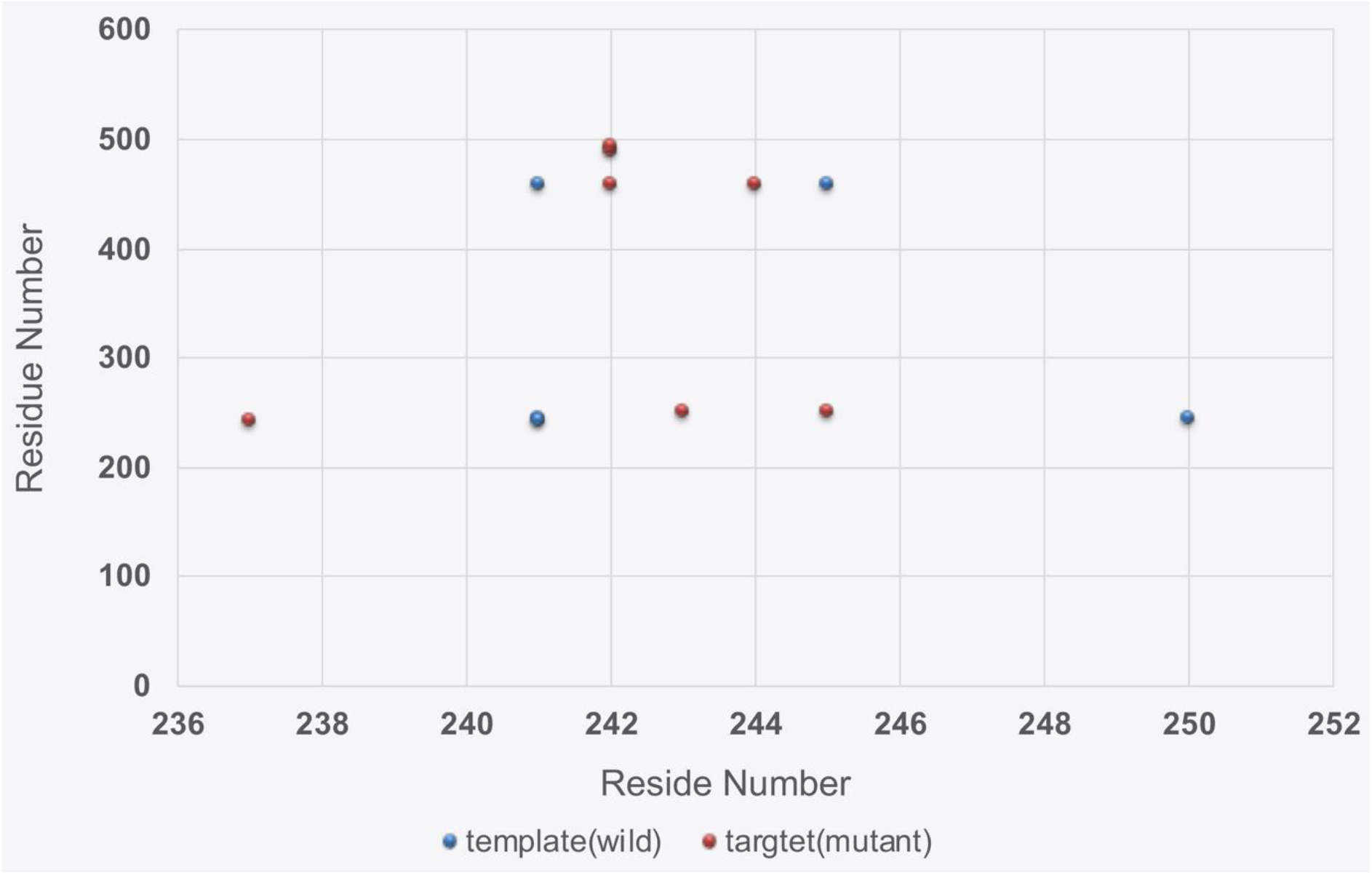
The differential contact map representing differentially stabilizing contacts. Red represents the contacts that are unique in the wild type and green represents those that are unique in the mutant form.

### 3.7. Protein-protein interaction (PPI) analysis

To determine the effects of the mutations on the NOD2-related downstream molecules and pathways, we assessed its interactions in a PPI network. We first constructed an extended network of the NOD2-related proteins based on their direct PPI neighbors and their interactions (Figure 8) and then the highly connected region from this network and their related pathways was extracted (Figure 8). The results identified Receptor-interacting serine/threonine-protein kinase 2 (RIPK2); autophagy related 16 like 1 (ATG16L1); caspase recruitment domain-containing protein 9 (CARD9); Tumor necrosis factor receptor-associated factor 6 (TRAF6); and inhibitor of Nuclear Factor Kappa B Kinase Regulatory Subunit Gamma (IKBKG) as the highly connected molecules with NOD2 that mainly are involved in apoptosis and immune-related pathways including mycobacteria (Figure 8).

**Figure 8.**
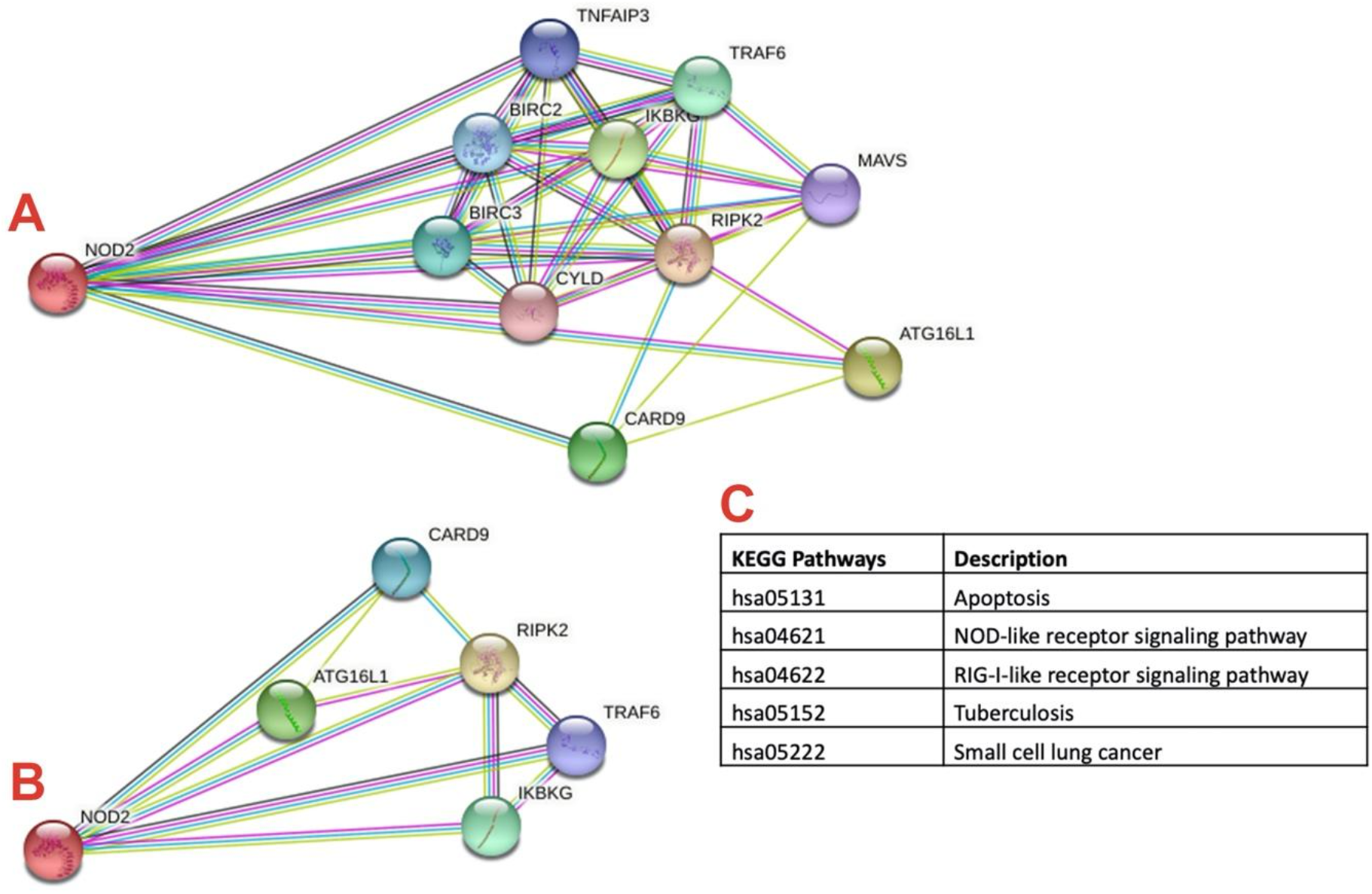
(A) The PPI network of the NOD2 receptor. (B) PPI networks were constructed, and the cluster contained the highly connected regions in the network identified by the Molecular COmplex DEtection plugin. (C) The related pathways of these highly connected proteins were identified and extracted. Nodes represent proteins and edges represent their interactions.

### 3.8. Prediction of post-translational modification sites

For further investigation of the impact of the point mutations, the prediction of post-translational modification sites was performed.

The NetPhos 3.1 and GPS 3.0 servers with 90% specificity predicted that the wild and mutant residues in both SNPs would not undergo phosphorylation or methylation.

Moreover, to see if the new lysine in the position 778 in the mutated protein could be targeted for ubiquitination, we used the BDM-PUB tool to predict ubiquitination sites. The results showed that this new residue introduces a new ubiquitination site in the mutated protein. (Supplementary Figure S1A).

Further analysis of the two SNPs on the PTM process by ProP 1.0 Server indicated that the new residues in position 908 and 778 introduce new propeptide cleavage sites in the mutant proteins that were not present in the wild type (Supplementary Figure S1B).

## 4. Discussion

In this study, we assess the impact of E778K and G908R on the structure and function of the NOD2 receptor. We explain the effect of the mutations and their role in the consequent behavior of the mutant NOD2 receptor. We showed that those mutations could impress the ligand recognition, stability of the protein, modulation of LRR conformation, and interaction with other proteins. Furthermore, the decreased flexibility in this region of those mutations could reduce the transformation of the inactive to active form of the protein and completely reduce susceptibility to the ligand, resulting in receptor loss of function.

Furthermore, these mutations introduced a new ubiquitination site and peptide cleavage sites in the protein that could impact the stability and half-life of NOD2.

Tanabe et al. conducted a mutational analysis of the NOD2 protein and reported that the G908R mutation causes loss of function (LOF) in NOD2^41^. However, they could not find the underlying mechanism for LOF.

The current study shows that G908R decreases the flexibility in the WHD–HD2–HD1 domain and so slows conversion to the active form of the receptor while increasing the rigidity in this region. Additionally, in the position of G908, the torsion angles in this region are unusual, and glycine is flexible enough to make these torsion angles. The presence of a larger R-side chain in this region forces the local backbone into an incorrect conformation and will disturb the local structure.

It has been shown that in the LRR–HD1 interface, the α3 helix (HD1) is packed against the α1 and α2 helices of the LRR, mainly through different interatomic contacts ^11^. E778 is located in α2 helices, and its interaction with its adjacent residues plays an important role in this interface. Our data showed that substitution of E to K in position 778 significantly changed the interatomic interactions in this region. We observed that this mutation decreased total flexibility of the protein, making it less capable of structurally converting to the active form.

Upon introducing those mutations in the LRR domain, the pattern of contacts between the residues in the CARD domain of NOD2 changes and may impress CARD-CARD interaction in the downstream reactions. The active site prediction results show that the G908R mutation significantly changed interatomic interactions, which contribute to changing the position and conformation ligand binding site. This may play an essential role in the MDP non-responsiveness of the mutant protein that was previously reported by in vitro experiments^41^. The mutations introduced a new ubiquitination site in the protein which may impress downstream interactions. Introducing new peptide cleavage sites upon mutation makes the mutant protein susceptible to proteases and reduces its half-life and stability.

The presence of E778K and G908R may not be the only determinant for infection development. Environmental factors and other transcriptional factors, including miRNAs, might play a role in poor outcomes. The presence of other mutations in the downstream molecules that may boost or compensate for the effects of these inborne errors is another possibility for lack of response to NTM.

This study helps to provide a comprehensive view of the potential role of inborne errors at E778K and G908R of NOD2 on the molecular mechanisms of diseases, potentially facilitating the development of Nontuberculous mycobacterial disease.

## Supplementary Materials

The following are available online at www.mdpi.com/xxx/s1, Figure S1: (A) Shows a new ubiquitination site in the E778K mutated protein. (B) Shows two new residues in position 908 and 778 of NOD2, introducing new propeptide cleavage sites.

## Ethics approval and consent to participate

Not applicable

## Consent for publication

Both authors agree with publication of final version.

## Authors’ contributions

MM conceived of the idea. SDA performed the calculations, collected the data, and performed data analysis. SDA and MM contributed to the interpretation of the results and provided critical feedback and helped shape the research, analysis, and manuscript. Both authors contributed to the final version of the manuscript.

## Acknowledgements

We thank Dr. Mallika Iyer for assistance with the analysis of contact maps and drawing the corresponding diagram.

## Availability of data and material

All data have been presented in the manuscript and supplemental file.

---

**Figure S1.**
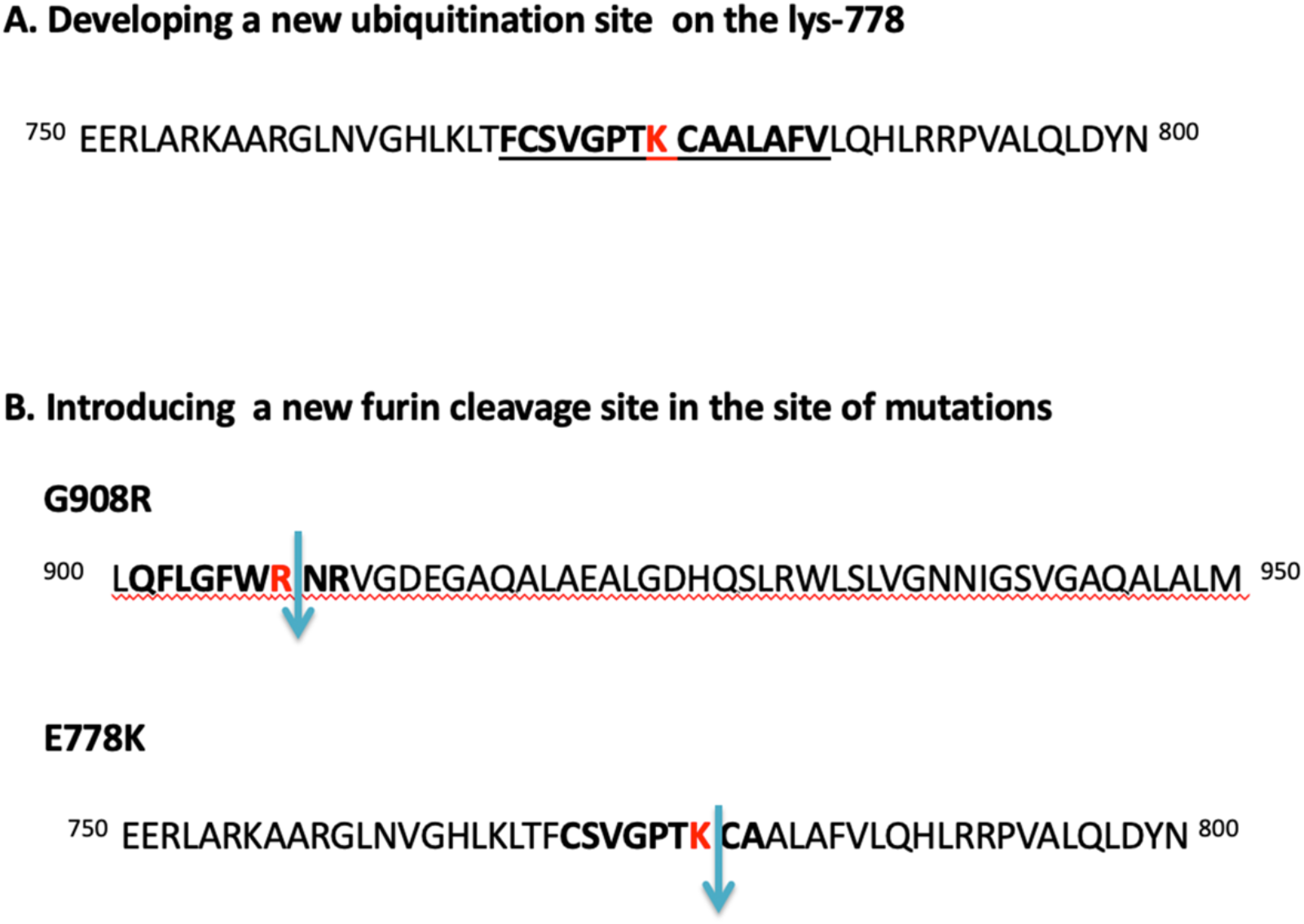
(A) Shows a new ubiquitination site in the E778K mutated protein. (B) Shows two new residues in position 908 and 778 of NOD2, introducing new propeptide cleavage sites.

